# Design and characterization of porous poly(glycerol-dodecanedioate) scaffolds for cartilage repair

**DOI:** 10.1101/2023.03.03.531009

**Authors:** Yue Qin, Sriharsha Ramaraju, Scott J. Hollister, Rhima M. Coleman

**Affiliations:** Department of Biomedical Engineering, University of Michigan, Ann Arbor, Michigan, United States of America; Wallace H. Coulter Department of Biomedical Engineering, Georgia Institute of Technology, Atlanta, Georgia, United States of America; Department of Mechanical Engineering, University of Michigan, Ann Arbor, Michigan, United States of America

**Keywords:** elastomer, porous scaffold, cartilage tissue engineering, ligand coating, parameter optimization

## Abstract

Synthetic polymeric scaffolds play an important role in establishing the microenvironment for chondrocytes in engineered cartilage. A three-dimensional pore network allows cell accommodation and supports extracellular matrix (ECM) production by chondrocytes. Ligand coating and biomechanical properties of scaffolds guide regeneration of functional cartilage by mediating cell attachment and establishing the local strain environment. Poly(glycerol-dodecanedioate) (PGD) is a novel biodegradable elastomer with nonlinear-elastic properties similar to native cartilage. However, its harsh curing environments limit the feasibility of common strategies for pore creation in polymeric scaffolds. Herein, we developed porous PGD (pPGD) scaffolds with tailorable pore structures using an inverse molding method and evaluated the range of scaffold structural parameters achievable and their subsequent mechanical properties. The influence of coating PGD with various ECM ligands on the cell shape, metabolic activity, and ECM production of human articular chondrocytes (hACs) was evaluated. pPGD scaffolds were created with pore sizes ranging from 250 – 1000 μm, resulting in 20 – 50% porosity. The morphology and metabolic activity of hACs on PGD were regulated by the type of ligand coating used. When compared to tissue culture plastic, PGD enhanced ECM production in monolayer cultures. Finite element analysis showed that the tensile strains that developed on the pores’ surfaces were at levels shown to be anabolic for hACs. The predicted strain profile varied with pore size and porosity under load, demonstrating that the pore structural parameters could be tuned to optimize cellular-level strains. These results suggest that pPGD scaffolds have the potential to guide cartilage regeneration.

**Statement of Significance:** Previous studies have established the importance of designing pore geometry and surface properties in engineered cartilage tissue constructs. This work reports the development and assessment of pPGD scaffolds with tunable pore and surface parameters for cartilage regeneration. The cellular-level strain that cells may experience inside the pores was influenced by the scaffolds’ pore geometry. Ligand coating on PGD balanced out the less ideal properties of the material itself and regulated the shape, attachment, metabolic activity, and ECM production of hACs during *in vitro* culture. These findings highlight how intelligent design of scaffold parameters can optimize chondrocyte function during 3D culture by tuning ligand presentation and cellular-level strain profiles.

## 1. Introduction

Articular cartilage has a limited ability to self-repair, which often causes traumatic injuries to progress into post-traumatic osteoarthritis (PTOA) [1]. Chondrocyte-based tissue engineering is an appealing approach for treating cartilage defects and preventing the onset of PTOA [2]. Among the biomaterials being investigated for cartilage tissue engineering, polymers are one of the most flourishing areas since they are easily processed into porous scaffolds that sustain cell proliferation and can be tailored for desired mechanical properties and degradation profiles [3]. An ideal polymer scaffold for articular cartilage regeneration should provide the following characteristics [4]: (i) a 3D interconnected porous structure to allow cell attachment, proliferation, differentiation, and extracellular matrix (ECM) production and promote nutrient and waste exchange, (ii) a biocompatible and bioresorbable substrate with controllable degradation rates, and (iii) mechanical properties similar to natural cartilage tissue to withstand the surrounding harsh biomechanical environment, to provide an optimally engineered 3D environment that reverses the chondrocyte dedifferentiation occurring during two-dimensional (2D) expansion and induces chondrocyte redifferentiation and maintains the chondrocyte phenotype.

Polymeric scaffolds have the potential to support chondrocyte function while protecting them from load. By incorporating strategies and techniques for creating, controlling, and characterizing scaffold architecture and cell-material interactions into the scaffold design process, the local environment can be tuned to enhance cell function. Modulations of scaffold parameters, such as pore size, total porosity, material surface chemistry, and biomechanical properties, have been shown to enhance cartilage tissue regeneration during *in vitro* co-culture [5]. Optimization of scaffold design via manipulation of these scaffold parameters to create functional cartilage normally starts with *in vitro* models and computational models [6-9]. The influence of these parameters on chondrocyte function have previously been demonstrated in citrate-based scaffolds [10-13]. Studies show proper pore size and porosity of a scaffold support the maintenance of the chondrocyte phenotype and promote the biosynthesis of cartilage-related ECM components [14-16]. These parameters not only determine the macroscale properties of neo-cartilage, which must facilitate joint function, but they also affect mechanotransduction in chondrocytes by altering the local environmental strain fields around the cells [17]. An optimal scaffold will conduct anabolic mechanical signals to the cells while protecting them from physiologic joint loads.

Scaffolds composed of ECM ligands create an environment that can preserve the normal phenotype of cells to promote the regeneration of cartilage-like constructs. Cell-matrix interactions play an important role in the maintenance of the chondrocyte phenotype. Chondrocytes can lose their original round shape in 2D settings such as large scaffold pores, leading to dedifferentiation and downregulation of collagen type II and aggrecan expression, which results in inferior tissue quality [18]. To mitigate this, ECM ligands that promote cell attachment and translate extracellular stimuli, such as strain, into intracellular signals [19-22], are incorporated into porous structures. For example, coating type I collagen (Col I) on scaffold surfaces can suppress morphological changes in chondrocytes in 2D cultures [23]. Type II collagen provides signaling cues via several receptors (e.g., integrin receptors α10β1 and α2β1) on chondrocytes to stabilize their differentiated morphology and promote ECM production [24, 25]. Hyaluronic acid (HyA), another important cartilage ECM ligand, can be recognized by chondrocytes via the surface receptor CD44 and provide positive influences on many chondrocyte pathways, including phenotypic regulation and ECM production [26]. While various hydrogels consisting of these ECM components do support the maintenance of the chondrocyte phenotype, they cannot fulfill the protective mechanical function of articular cartilage [27]. Biodegradable polyesters are usually mechanically tough, but their hydrophobicity leads to inefficient cell attachment, requiring surface modification to enhance their cell affinity [28].

Cells can sense the external mechanical and ligand cues and translate those cues into intracellular signaling, which influences cell behavior. There are many cellular structures and pathways involved in mechanical sensing and transduction into changes in gene expression and protein production. These include structural elements, such as stress fibers, focal adhesions, and integrins, activating mechanotransduction pathways such as RhoA/ROCK and Yap/Taz [17]. Binding to ECM ligands induces mechanical sensing through actin fibers and activation of integrins, cell subsequently adjusts its cell-ECM interaction strength and finally alters its shape, proliferation, migration, and phenotype through multiple pathways [17, 29, 30]. The scaffold design process hence requires the optimal structural parameters and ligand coating for the desired level of mechanotransduction since chondrocytes reorganize the ECM components of cartilage in response to the external load. Previous work has shown that, in monolayer culture, loading between 3–10% cyclic tensile strain and 0.17–0.5 Hz led to anabolic responses of chondrocytes, and beyond that range caused catabolic events to predominate [31]. Due to the 2D nature of the pore surface inside the scaffold for cell attachment, this important design parameter should be incorporated into the design of polymer scaffolds.

Synthetic polymers such as polylactic acid (PLA), polyglycolic acid (PGA), their copolymer polylactic-co-glycolic acid (PLGA), and polycaprolactone (PCL) have been used as scaffolds for years in the field of cartilage tissue engineering. However, these synthetic polymers exhibit linear elastic behavior and a mismatch of mechanical properties with native cartilage tissue, leading to device failure [32, 33]. Biodegradable elastomers like polyglycerol sebacate (PGS) and poly (1,8 octanediol-co-citrate) (POC) have been increasingly studied for soft tissue repair applications [13, 34, 35]. A study has shown that POC with nonlinear elasticity showed higher sGAG production and lower hypertrophy after 4 weeks of *in vitro* cell culture compared to PCL, suggesting that rubber-like biodegradable polyester elastomers have the ability to support chondrocyte redifferentiation and could facilitate its function [13].

The purpose of this paper is to describe the design and characterization of 3D porous scaffolds made of poly(glycerol-dodecanedioate) (PGD) for cartilage tissue engineering. PGD is a novel biodegradable elastomer formed by the polycondensation of glycerol and dodecanedioic acid [36]. It was first reported in 2010 for potential use in soft tissue engineering applications because of its elastomeric mechanical properties, low *in vitro* degradation rate, shape memory behavior, and good biocompatibility [36-38]. PGD has nonlinear elastic properties similar to those of various soft tissues at body temperature and elastic-plastic behavior at room temperature [36, 37]. Moreover, the mechanical properties of PGD can be controlled by varying the curing time and temperature, resulting in materials with a range of moduli comparable to soft tissues, e.g., cartilage [37]. Those properties make PGD a viable model choice for scaffold optimization in cartilage tissue engineering. However, the application of PGD for cartilage tissue engineering is hindered by the harsh curing conditions (high temperature and vacuum), which limit the number of strategies that can be used to create a porous structure while maintaining the stiffness of the scaffold. Herein, an inverse molding method was used to create porous PGD (pPGD) scaffolds with tailorable, interconnected pore structures. The ranges of achievable scaffold structural parameters (pore size and porosity) were evaluated with microcomputed tomography (micro-CT), and their mechanical properties were evaluated via compressive testing. The effects of ligand coating on the morphology, metabolic activity, and ECM synthesis of human articular chondrocytes (hACs) were also evaluated. Finally, the effect of porosity and pore size on the local strain fields inside simulated pPGD scaffolds was evaluated using finite element analysis.

## 2. Materials and methods

### 2.1. PGD fabrication

#### 2.1.1. Prepolymer and curing

PGD prepolymer was synthesized and then cured following the methods described by Solorio et al. [37]. Briefly, PGD pre-polymer was synthesized by mixing glycerol and dodecanedioic acid in a 1:1 molar ratio at 120 °C under nitrogen and stirring conditions for 24 h. The viscous pre-polymer was then cast into silicone molds and transferred to a vacuum oven at 130 °C for 48 h. A vacuum was pulled and maintained at 90 mTorr for the duration of the curing process, to get the solid, nonporous PGD blocks.

#### 2.1.2. Porous PGD scaffold fabrication

The pore size of the pPGD scaffold was controlled using Teflon-coated stainless-steel wires, as outlined in **Figure 1a**. Due to their chemical inertness and thermostability, Teflon-coated wires were easily removed after PGD curing and therefore presented a good choice for creating voids in the thermoset PGD scaffolds. PGD prepolymer was synthesized as described above. To control the wire position, polylactic acid (PLA) guides were designed in SOLIDWORKS software and then printed on a 3D printer (Creator Pro, FlashForge, China). The PLA guide was placed into a silicone mold, and Teflon-coated stainless-steel wires (McMaster-Carr, USA) of 250 μm, 500 μm, 750 μm, or 1000 μm in diameter were pierced through the wall of the silicone mold when their final position was located by the PLA guide (**Figure 1a**). Porosity was controlled by decreasing the spacing between wires, thereby increasing the wire density. The PGD prepolymer was then cast into the mold and cured under a 90 mTorr vacuum maintained at 130 °C for 48 h. After curing, the wires were removed, and the pPGD scaffolds were peeled out of the silicon mold. Each pPGD scaffold was then cut into cylinders 6 mm in diameter and 3 mm in height.

**Figure 1.**
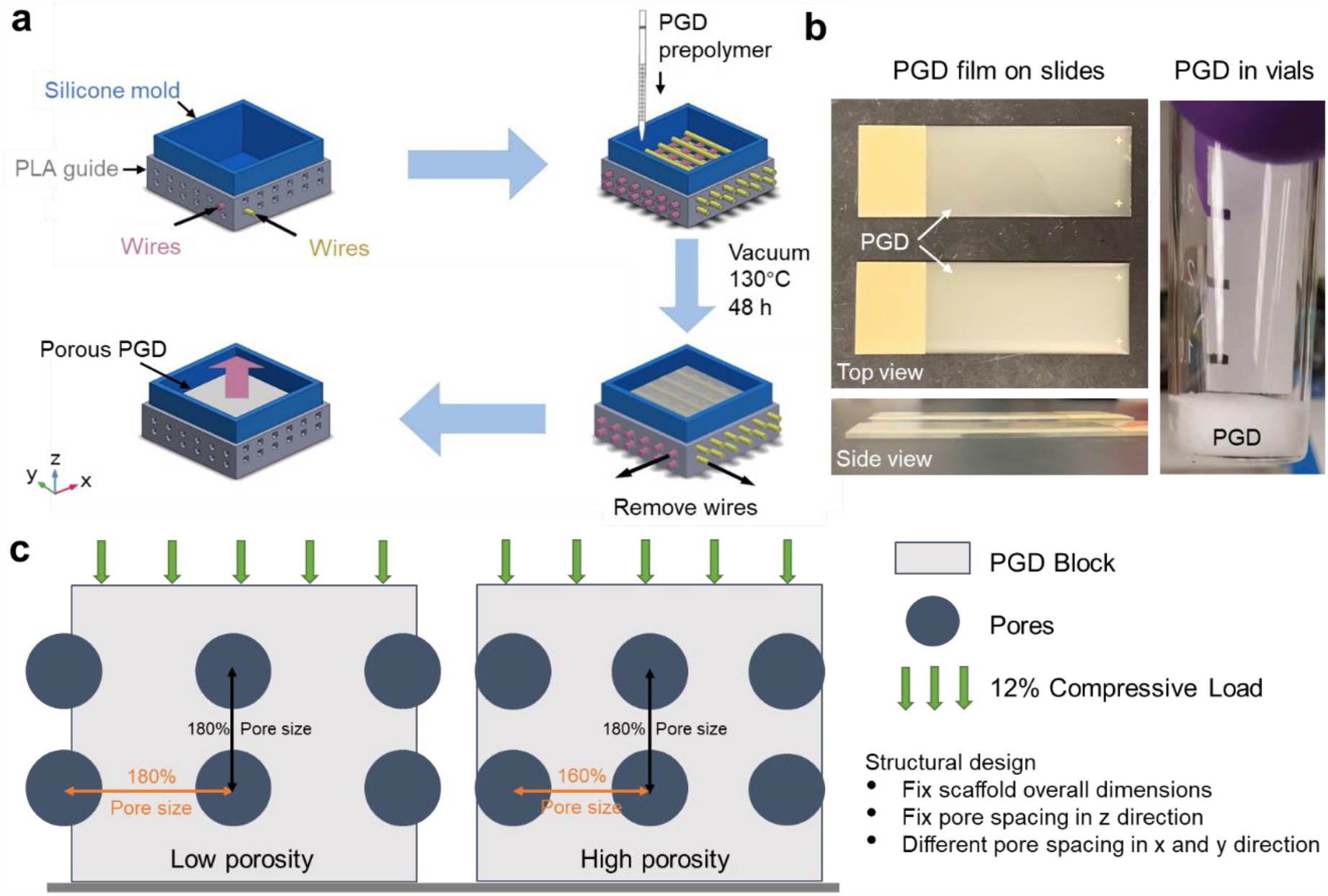
Design of pPGD scaffolds and PGD films. (a) Schematics of pPGD scaffolds fabrication via inverse molding. The silicone mold and Teflon coated stainless steel wires were assembled as the inverse mold. Wires were aligned along x or y directions and organized alternately (layer-by-layer) in z direction. The position of wires was precisely controlled by polylactic acid (PLA) guides. (b) Digital images of PGD films after curing. Flat surface of PGD were created as PGD film on slides or in glass vials. (c) Schematics of geometries design for simulated pPGD scaffolds. The porosity of simulated pPGD scaffold was altered by changing the pore spacing horizontally.

#### 2.1.3. PGD surface fabrication and preparation

Two-dimensional (2D) PGD surfaces were used to determine the morphology, metabolic activity, and ECM production of hACs in the monolayer settings. PGD prepolymer was synthesized as described above before being cast onto positively charged microscope slides (InkJet Plus Microscope Slides, Fisherbrand, USA) or 3 mL glass vials (Restek Corporation, USA). The PGD prepolymer spontaneously spread to a flat film on the positively charged slides or formed a flat surface on the bottom of glass vials (**Figure 1b**), and was then cured with the same conditions as previously described (90 mTorr vacuum, 130 °C, 48 h). All the PGD surfaces were soaked in the low glucose Dulbecco’s modified Eagle’s medium (DMEM) (Gibco, Biosciences, Dublin, Ireland) for 7 days with three media changes to remove any possible cytotoxic byproducts that could dissolve in the media.

### 2.2. Scaffold structural parameters analysis

The scaffold micro-geometry was reconstructed from microcomputed tomographic (μCT) images acquired using Scanco μCT 100 system (SCANCO Medical, Switzerland). Samples were scanned using a 4 μm resolution. The 3D models of the scaffold geometry were reconstructed from the μCT images using Materialise Mimics software, and the scaffold volume and total volume were quantified throughout the sample geometry. The porosity was calculated using the following equation after quantifying the scaffold volume.

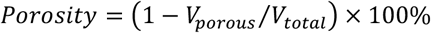

Where *V*_*porous*_is the volume of the porous structure and *V*_*total*_is the total volume of the enclosed structure.

### 2.3. Mechanical testing

To analyze the effects of pore parameters on the mechanical properties of pPGD scaffolds, compression testing was conducted at 37 °C. MATLAB (The MathWorks Inc., USA) software was used to fit a one-term Ogden model to experimental data to obtain the shear modulus of each pPGD scaffold. Compressive testing was conducted on 6 mm pPGD cylinders within a custom-built temperature control chamber using an MTS system equipped with a 500 N load cell and a metal platen. Samples were tested in the chamber maintained at body temperature (37 °C). Compression was applied at a rate of 2 mm/min, and samples were compressed to about 60% strain at 37 °C. The mechanical test data obtained at 37 °C was then fitted to a one-term Ogden constitutive model for nonlinear hyperelastic materials using custom MATLAB algorithms. In brief, the one-term Ogden model was based on a strain energy function of the following form.

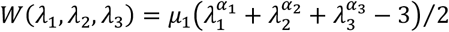

Where W was the strain energy function, *λ*_*i*_ denoted the stretch ratios in the *x*_*1*_, *x*_*2*_, and *x*_*3*_ directions, and *μ*_1_ and *α*_*i*_ were constants that were fit to the experimental data. For the uniaxial compression test in this study, assuming the specimen was tested in the *x*_*3*_ direction, the resulting normal 1st Piola-Kirchoff stress was calculated as the following equation.

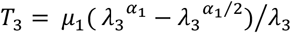

The normal stress was calculated from the experimental compressive test by dividing the applied force by the initial specimen area. The model coefficients (*μ*_1_ and *α*_1_) were then fitted to the experimental stress using the *fmincon* routine in the MATLAB optimization toolbox. The results were further constrained using the Baker-Eriksen inequality shown below [37].

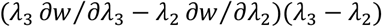

The coefficient of determination (R^2^) was calculated for fitting quality, and the estimated shear modulus (*μ*) of the specimen was calculated by the following equation.

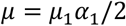

### 2.4. Surface coating and characterization

PGD surfaces were sterilized with sterile 70% ethanol through a 30-minute ultrasonic wash. PGD surfaces were coated with the ECM ligands collagen type I (Col I) or hyaluronic acid (HyA), via physical absorption. Briefly, HyA powder (molecular weight ∼1.5 mDa, Lifecore, USA) was dissolved in a sterile phosphate-buffered saline (PBS) to achieve a stock concentration of 0.1 % (w/v). Col I from rat tail tendon dissolved in 0.1 M acetic acid solution (Corning, NY, USA) was purchased for a stock concentration of 0.1 % (w/v). The aqueous solution of HyA or Col I (1 ml) was pipetted onto the PGD surfaces in slide dishes (Nunc rectangular dish, Thermo Scientific, USA) or in the glass vials, until the PGD surfaces were fully immersed. All samples were air-dried for 2 days in a sterile environment at room temperature to prepare the coated substrates.

The apparent water contact angles of the TCP and all the PGD samples were tested by a DSA100 type contact angle meter using a static sessile drop method. The water contact angle was measured by depositing 5 μl of an ultrapure water droplet on each sample surface. The angles of water droplets on the material surfaces were determined immediately after the deposition of droplets.

### 2.5. Cell Seeding and Culture Conditions

Human articular chondrocytes (hACs) from a healthy young male donor (age 19, CELLvo, StemBioSys, USA) were expanded in the growth media: Low glucose DMEM (Gibco, Biosciences, Dublin, Ireland) supplemented with 10% v/v FBS, 100 U/ml penicillin and 100 μg/ml streptomycin, 1 ng/ml TGF-β1, 10 ng/ml PDGF-BB, and 5 ng/ml FGF-2, until the plate reached confluency. hACs were seeded with a seeding density of 1×10^6^ cells/cm^2^ onto the top surface of coated PGD on the slide, 3 mL glass vials, or 12-well plates (tissue culture plastic, control group) and then incubated for 3 h for cell subsidence. After cell subsidence, the growth media was replaced with redifferentiation media, which was formulated as high glucose DMEM (Gibco, Biosciences, Dublin, Ireland), 1.25 mg/ml bovine serum albumin, 100 U/ml penicillin and 100 μg/ml streptomycin, 1% v/v ITS + Premix (Corning, NY, USA), 10 ng/ml TGF-β1, 40 μg/ml L-Proline, 50 μg/mL ascorbic acid 2-phosphate, 1 mM sodium pyruvate, and 100 nM dexamethasone. The hAC-seeded PGD and hACs in the 12-well plates were then cultured in the humidified incubator (37 °C, 5% CO_2_) for 2 or 28 days.

### 2.6. Cell attachment analysis

To analyze the effects of ECM ligand coating on cell attachment and cell shape, hACs were seeded onto tissue culture plastic (TCP) and PGD surfaces coated with 0.1% w/v collagen type I (PGD-Col I), 0.1% w/v hyaluronic acid (PGD-HyA), or without coating, and then cultured for 2 days as described above. The constructs were then examined by F-actin staining using rhodamine phalloidin (Invitrogen, USA) following the manufacturer’s instructions, and images of hACs were taken from the fluorescence microscope or Nikon A1 Confocal. Briefly, the samples were fixed with 10% neutral buffered formalin for 30 min and then permeabilized with 0.1% v/v Triton X-100 for 5 min. The samples were strained with a 300 nM phalloidin solution for 30 min at room temperature. To assess cell attachment, the perimeter and area of each human articular chondrocyte were quantified by analysis of the images via ImageJ 1.53c software. The following equation calculated the circularity of each cell.

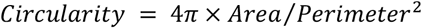

### 2.7. Metabolic activity analysis

Chondrocyte metabolic activity was analyzed via Cell Counting Kit-8 (CCK-8, Dojindo, Japan) following the manufacturer’s instructions. Briefly, hACs were seeded on PGD in vials or TCP before being cultured for 28 days. For each test, 10% v/v CCK-8 solution in high glucose DMEM was added to each vial on days 2, 5, 8, 12, 15, 20, 25, and 28, and the vials were incubated at 37 °C for 4 h. The absorbance of the solution in each vial was measured at 450 nm.

### 2.8. Biochemical analysis

To analyze the effect of ECM ligand coating on sulfated glycosaminoglycan (sGAG) production, a 1,9-dimethyl methylene blue (DMMB) assay was conducted on hACs in TCP or hAC-seeded PGD in vials after a 28-day culture as previously described [39, 40]. Briefly, the matrix generated on the top surface of PGD or TCP during the culture was digested with papain [39]. The absorbance was measured at 525 nm and 595 nm and compared to chondroitin sulfate standards [40].

### 2.9. Finite element analysis

To determine how pore size and porosity affect the strains that develop within the pores of pPGD scaffolds during loading, finite element analysis (FEA) was performed on the simulated geometries of pPGD scaffolds using COMSOL. The simulated pPGD geometries were modeled as 3 mm cubes with pore sizes of 250 μm, 500 μm, 750 μm, or 1000 μm. To vary the porosity of the model, the distance between the centers of each pore was fixed at 160% pore size for high porosity groups (High or H) or 180% pore size for low porosity groups (Low or L) (**Figure 1c**). All models were meshed with a tetrahedral volume element with a similar mesh size (minimum mesh size = 0.055 mm). The one-term Ogden constants of nonporous PGD that were obtained from mechanical tests and hyperelastic fitting as described above were applied to the scaffold mesh elements. The lower surface of the model was constrained in all directions, and a 12% prescribed displacement (0.3 mm) in the z-direction was applied to the upper surface. This boundary condition was chosen to mimic the level of cartilage deformation in the knee joint during 30 min of standing, based on the highest mean strain (12%) presented over the femoral-tibial contact area [41]. The first principal strain (maximum tensile strain) was calculated in the central pores of each simulation. The percentages of nodes on the central pore surface that denoted a strain range of 3 – 10 %, which has been shown to be beneficial for chondrocyte function [31], were calculated as %Beneficial strain.

### 2.10. Statistical analysis

Unless indicated otherwise, results were analyzed using a one-way ANOVA and Tukey post-hoc test for multiple comparisons in GraphPad Prism 8.0 (GraphPad Software, San Diego, CA). The porosity, surface area, and shear modulus of pPGD scaffolds were compared across different pore size groups. The cell area, circularity, and sGAG production of hAC cultured on PGD were compared across different coating groups. The metabolic activity of hAC cultured on PGD was compared across different coating groups on the same day of culture. The criterion for statistical significance was p < 0.05 in all tests. Unless indicated otherwise, results are expressed as mean ± standard deviation (SD). Sample size (n) is indicated in the corresponding figure legends.

## 3. Results

### 3.1. Geometry parameters of porous PGD scaffolds fabricated by inverse molding

The 3D reconstructions of the pPGD μCT images confirmed that the inverse molding approach successfully fabricated PGD scaffolds with the desired pore diameters, well-aligned pore structure, and good interconnectivity (**Figure 2a**). The porosity of pPGD scaffolds was successfully controlled by changing the diameter of the wires. Increasing the pore diameter while maintaining the distance between each wire resulted in increased porosity, except for 1000 group (**Figure 2b**). There was no statistically significant difference between the 1000 group and the 500 or 750 group since the designed distance between each wire in the pPGD scaffold with 1000 μm pore impaired the number of pore channels per structure. Pore sizes of pPGD scaffolds achieved a range of 250 - 1000 μm, resulting in porosity in the range of 20 – 50%. The creation of a pore network inside bulk PGD significantly increased the surface area of the constructs (**Figure 2c**). Changing the pore size above 500 μm led to no significant change in the total surface area of the pPGD scaffold, while the 250 μm group showed a higher surface area than other porous groups.

**Figure 2.**
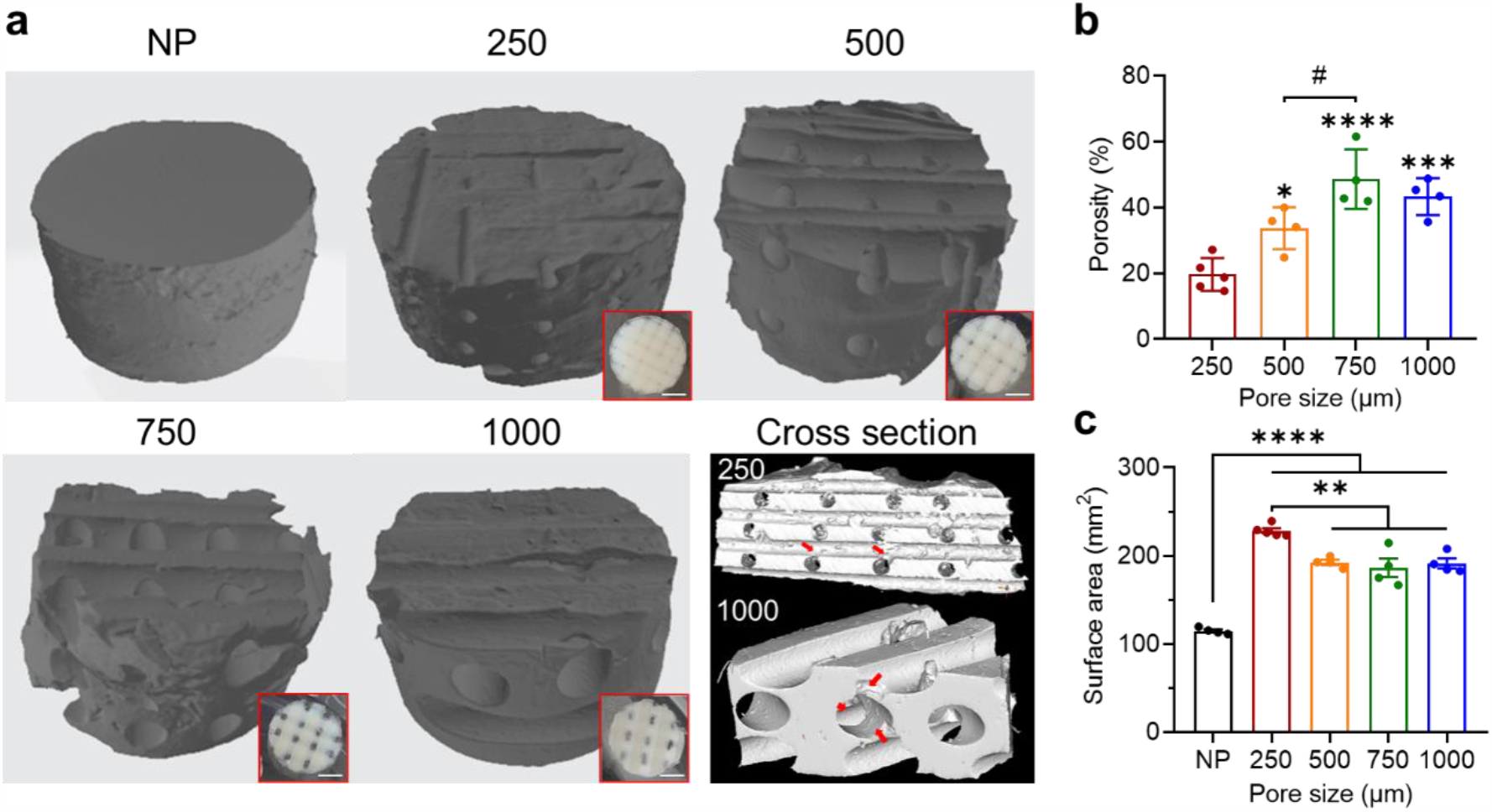
Geometries of porous PGD scaffolds. (a) 3D μCT image of nonporous PGD block and pPGD scaffolds with varying pore diameters (250 μm, 500 μm, 750 μm, or 1000 μm). NP denotes non-porous PGD scaffold. The digital pictures (top view) in red boxes showed pPGD scaffolds immersed in 70% ethanol. Scale bar: 2 mm. (b) Quantification of the porosity of pPGD scaffold with varying pore diameters. n = 4 scaffolds per group. Significant differences between 250 and other groups are indicated by * p < 0.05, *** p < 0.001, **** p < 0.0001. Significant difference between 500 and 750 is indicated by # p < 0.05. (c) Quantification of the total surface area of pPGD scaffold with varying pore diameter. n = 4 scaffolds per group. Data represented as mean ± s.e.m. Significant differences are indicated by ** p < 0.01 and **** p < 0.0001.

### 3.2. Geometry parameters of porous PGD scaffolds influenced mechanical properties

The stress-strain curve showed all groups of pPGD scaffolds and nonporous PGD bulk had nonlinear behaviors during compression (**Figure 3a**). The R^2^ value (> 0.95) indicated that the one-term Ogden model fit all the pPGD scaffolds well. (**Table 1 & Figure 3a**). The scaffold with higher porosity had less stiff nonlinear elastic behavior (**Figure 2b & 3b**). The shear modulus of the whole pPGD scaffold decreased with increasing porosity (**Figure 3c**). The pore structure had an influence on the maximum stretch ratio and the shear modulus among all the scaffolds, indicating the stiffness and mechanical behavior of pPGD scaffolds under compressive load can be altered by changing the pore parameters.

**Table 1.** Constants (μ1, α1) and R^2^ values of one-term Ogden model fitting PGD mechanical test data at 37°C. Results presented as mean ± standard deviation.

|  | NP | 250 | 500 | 750 | 1000 |
| --- | --- | --- | --- | --- | --- |
| $\mu_1$ | $0.631 \pm 0.108$ | $0.519 \pm 0.077$ | $0.466 \pm 0.036$ | $0.226 \pm 0.085$ | $0.240 \pm 0.033$ |
| $\alpha_1$ | $1.706 \pm 0.354$ | $1.327 \pm 0.279$ | $1.488 \pm 0.146$ | $1.197 \pm 0.499$ | $1.602 \pm 0.370$ |
| $R^2$ | $0.990 \pm 0.0063$ | $0.988 \pm 0.0064$ | $0.955 \pm 0.0160$ | $0.996 \pm 0.0044$ | $0.989 \pm 0.0108$ |

**Figure 3.**
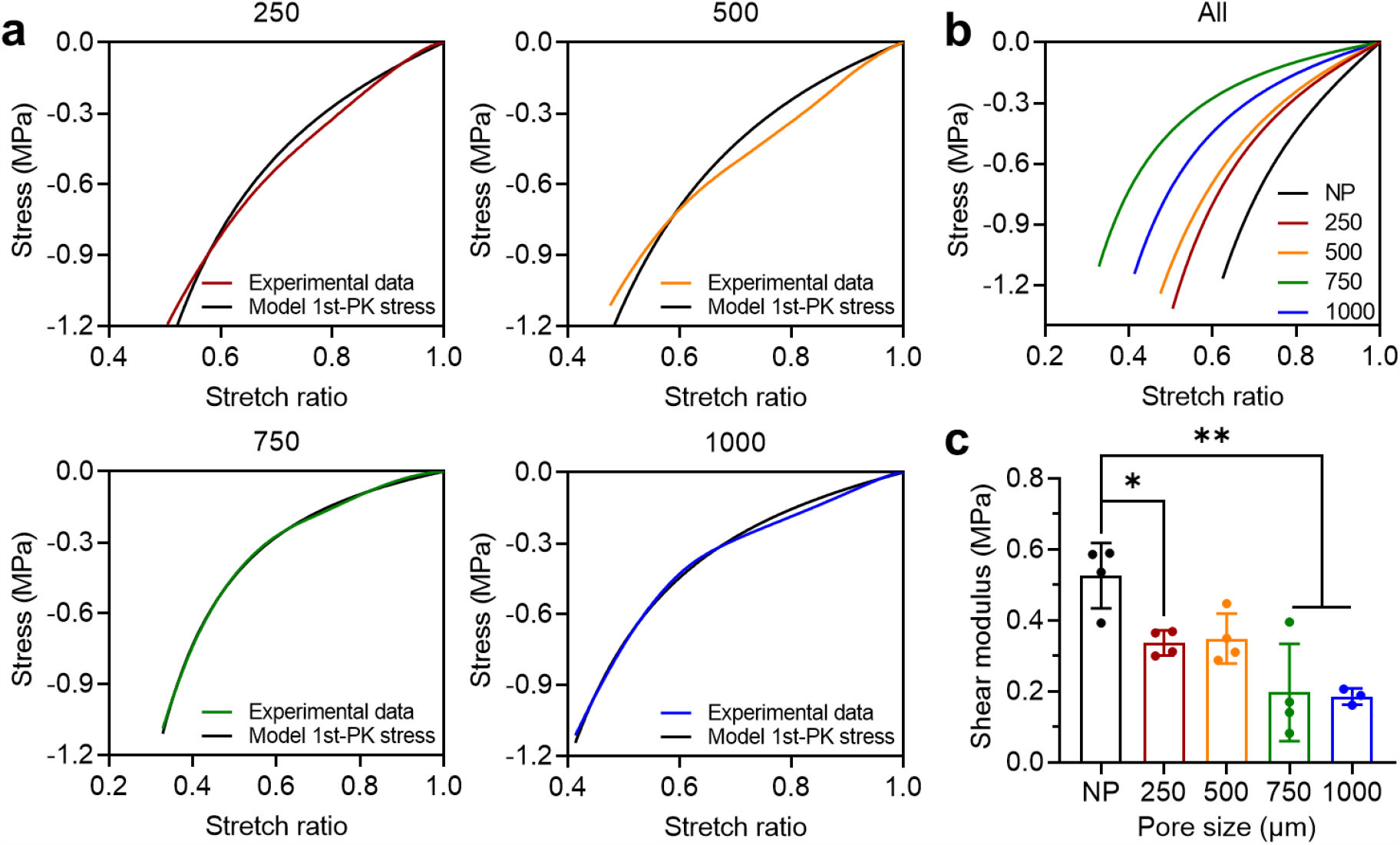
Pore size influenced the mechanical properties of pPGD scaffolds. (a) Stress-strain curves of pPGD scaffolds with varying pore sizes during compressive testing and corresponding one-term Ogden fitting curves. (b) Ogden fitting curves of all the pPGD scaffold groups and nonporous PGD. (b) Shear modulus of pPGD scaffolds with varying pore diameters. n = 4 scaffolds per group. Significant differences are indicated by * p<0.05 and ** p<0.01.

### 3.3. ECM ligand coatings influence chondrocyte function on PGD

The surface hydrophilicity of various PGD surfaces was comparatively evaluated by a water contact angle measurement using the sessile drop method. The water contact angle determined on the pure PGD surface was approximately 63°, which was similar to tissue culture plastic (TCP). The water contact angle of pure PGD was significantly greater than that of Col I or HyA coated surfaces, indicating that ligand coating increased the hydrophilicity of PGD surfaces and could improve cell affinity (**Figure 4a**). When PGD was coated with collagen type I rather than hyaluronic acid, the surface hydrophilicity was noticeably higher (**Figure 4a**).

**Figure 4.**
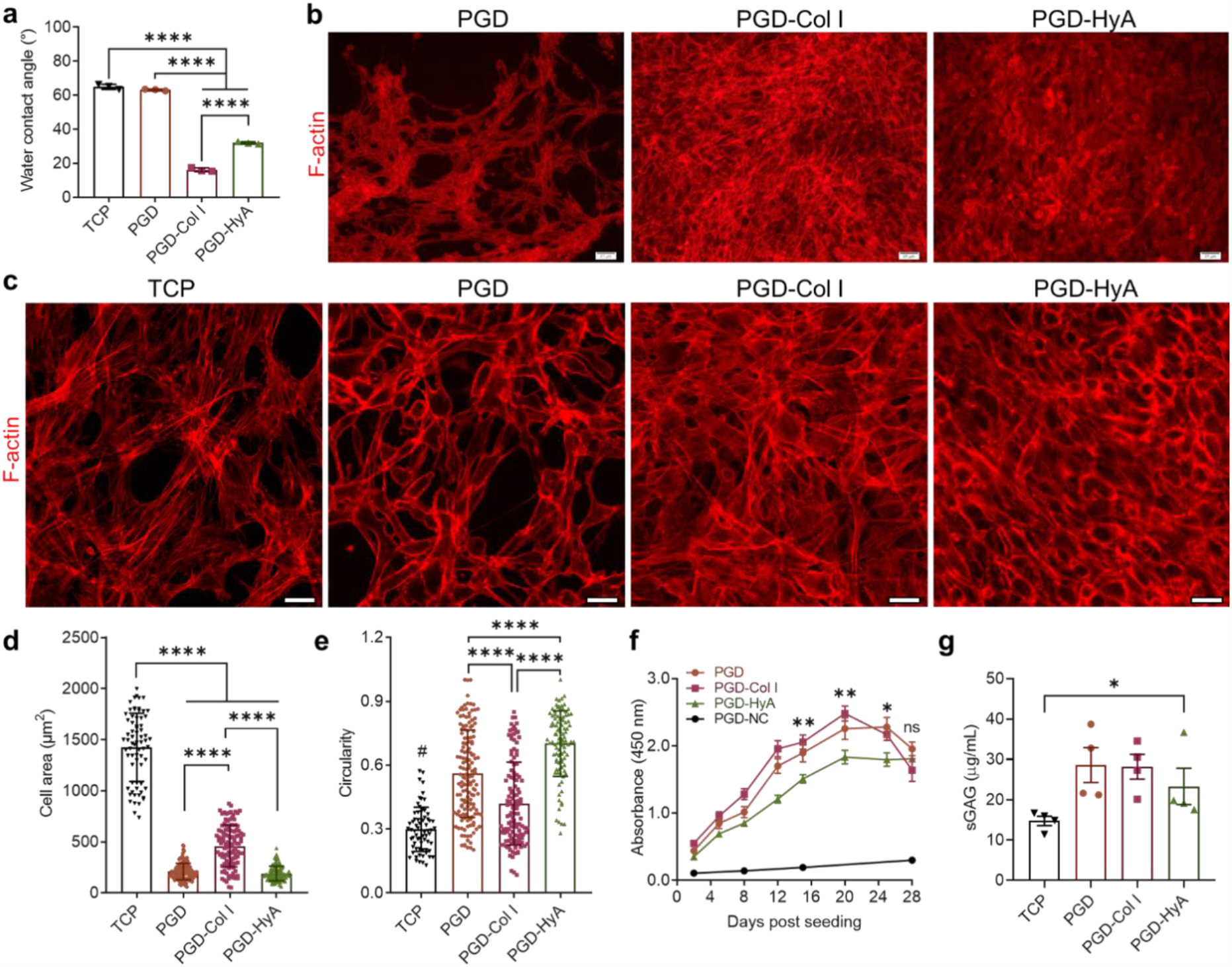
ECM ligands composition influenced chondrocyte behavior on PGD. (a) Water contact angles of PGD surface coated with different ligands. n = 3. Significant differences are indicated by **** p < 0.0001. (b) Fluorescence images of phalloidin TRITC staining (F-actin, red) on chondrocytes-seeded PGD films with no coating (PGD), Col I coating (PGD-Col I) or HyA coating (PGD-HyA) after 2 days of culture. Scale bar: 20 μm. (b) Confocal images of phalloidin TRITC staining (F-actin, red) on chondrocyte-seeded PGD films with different coating after 2 days of culture. TCP denotes tissue culture plastic. Scale bar: 20 μm. (d) Quantification of the area of each hAC on PGD. Significant differences are indicated by **** p < 0.0001. (e) Quantification of the circularity of each hAC on PGD. Significant differences among PGD, PGD-Col I, and PGD-HyA are indicated by **** p < 0.0001. Significant differences between TCP and other groups are indicated by # p < 0.0001. (f) Metabolic activities of hACs on PGD coated with different ligands in 28-day culture. PGD-NC denotes PGD with no cell seeded. n = 8 constructs per group. Significant differences among PGD, PGD-Col I, and PGD-HyA on the same day of culture are indicated by * p < 0.05 and ** p < 0.01. Data represented as mean ± s.e.m. (g) sGAG production of hACs on PGD coated with different ligands after 28 days of culture. n = 4 constructs per group. Significant difference between PGD and TCP is indicated by * p < 0.05. Data represented as mean ± s.e.m.

F-actin staining was conducted to evaluate the influence of ligand coating on cell attachment and cell shape after 2 days of culture. Human articular chondrocytes (hACs) attached and proliferated on PGD surfaces, demonstrating the high degree of cytocompatibility of PGD. (**Figure 4b**). Overall, the fluorescence staining of F-actin showed the cell number per area was lower on non-coated PGD films than the coated ones, and cell morphology was altered by the ligands. While the PGD alone and TCP groups accommodated fewer cells with voids between cell clusters forming a honeycomb-like structure, the PGD-Col I and PGD-HyA groups had much more cells with 100% confluency and dense, multi-layered structures of cells (**Figure 4b & 4c**). This suggested that ligand coatings improved hAC adhesion and proliferation on PGD. The type of ligand coating also influenced the cell morphology of hAC on PGD. In comparison to the PGD-Col I group, the HyA-coated PGD retained more hACs with round cell shape (**Figure 4c**). hAC grown on collagen type I surfaces exhibited a flat, stretched polygonal shape with an abundance of stress fibers. In contract, when grown on the PGD-HyA, cells were small and spherical, and the actin was distributed evenly beneath the cell membrane, which is associated with maintenance of chondrocyte phenotype [42, 43]. The quantification of cell area and circularity through ImageJ analysis on Day 2 supported this (**Figure 4d & 4e**). Col I presentation on the PGD surface increased hAC spreading with the largest surface hydrophilicity and single cell area, and the lowest circularity, while HyA coating maintained the smaller cell and rounder cell shape on PGD that is typically seen in 3D culture. Furthermore, despite the influence of ligands on cell behavior, PGD alone improved the maintenance of the chondrocyte phenotype in two-dimensional (2D) culture. Chondrocytes grown on TCP on day 2 adopted an increased fibroblast-like morphology compared to the other PGD groups (**Figure 4b - 4e**). Even though the pure PGD surface had significantly lower hydrophilicity than coated groups and similar hydrophilicity to TCP, it retained a smaller cell area, a rounder cell shape, and a higher cell circularity than TCP and PGD-Col (**Figure 4a - 4e**).

The effect of ECM ligand coating on hAC’s metabolic activities and ECM production was then investigated in 28-day cultures. The metabolic activity of hACs increased in all groups during the culture (**Figure 4e**), indicating their good cytocompatibility and support for cell growth. The metabolic activity of hACs varied depending on the PGD surface (**Figure 4e**). The metabolic activity of hACs cultured on Col I-coated PGD films was the highest, while that of those cultured on HyA-coated PGD films was the lowest; however, this changed after day 20. From day 20 to day 28, hACs cultured on PGD-Col I had gradually decreased metabolic activity, whereas hACs cultured on PGD-HyA maintained their metabolic activity level. From day 20 to day 24, hACs cultured on uncoated PGD had a stable level of metabolic activity, which then decreased by day 28. On day 28, there were no significant differences in the metabolic activity between the PGD, PGD-Col I, and PGD-HyA groups. PGD controls with no cells (PGD-NC) exhibited an absorbance much lower than other groups throughout the culture period, suggesting that PGD degradation byproducts had no effect on the accuracy of the metabolic assay. All the PGD groups supported chondrocyte anabolic function. Chondrocytes cultured on PGD produced more sGAG on day 28 than those on TCP, and uncoated, Col-coated, and HyA coated PGD generated the same amount of sGAG (**Figure 4f**).

### 3.4. Cellular-level pore strain fields can be tuned in porous PGD scaffolds

Finite element analysis was used to determine the effects of pore structure on cell-level strains that develop inside the pores on the pPGD scaffolds. We focused our analysis on the central pore in each model to reflect the maximum tensile strains (first principal strains) that develop on the pore surfaces. In all models, the majority of the central pore’s surface area displayed low surface strains (0 – 10%), and their overall strain fields were comparable (**Figure 5a**). The intersection area between the top and bottom pores experienced the lowest strain. As a result, the top and bottom sides of each central pore displayed a uniform distribution of the low-level strain, while the left and right sides of each central pore displayed a pattern of alternate high and low strain. Among all the groups, the 1000 μm pore size group had the highest tensile strains (> 16%) on the central pore’s intersecting area (**Figure 5a**). This was anticipated given that the scaffold’s bulk stiffness and matrix volume both decreased as pore size and porosity increased, and the scaffold with larger pores had more stress concentration regions than the one with smaller pores.

**Figure 5.**
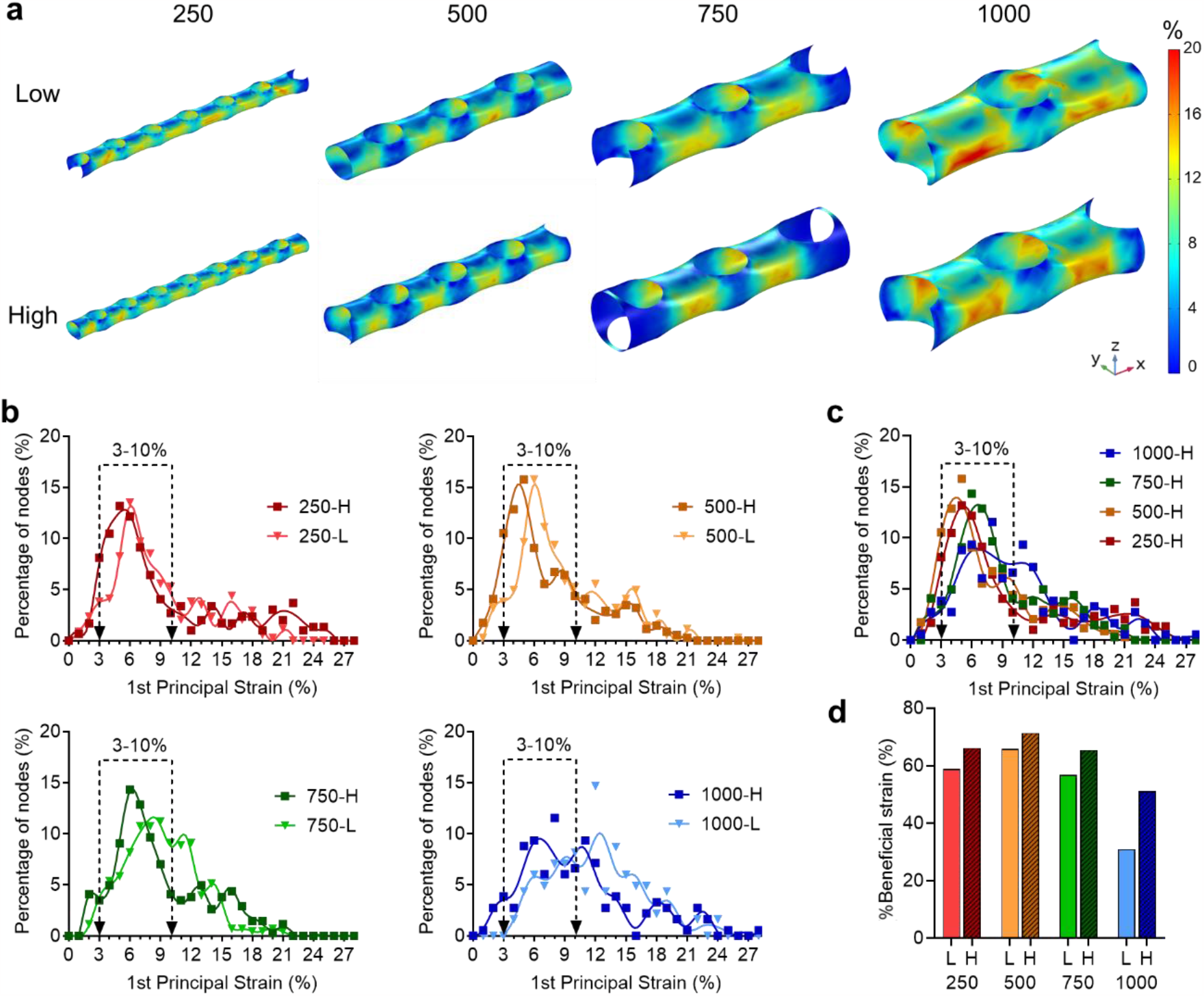
Finite element analysis of strain fields in pPGD scaffolds. (a) First principal strain fields of simulated pPGD models with 250 - 1000 μm pore size and two levels of porosities. “Low” denotes low porosity, “High” denotes high porosity. (b) The distribution curves of first principal strain on the central pore of the pPGD models. “L” denotes low porosity, “H” denotes high porosity. y-axis: the percentage of nodes on the central pore that experienced a certain strain. Dotted line: the specific period of the distribution curve that experienced the tensile strain that shown to be beneficial to chondrocyte. (c) The distribution curves of first principal strain on the central pore of all the high porosity groups of pPGD models. (d) Quantification of %Beneficial strain on the central pore of pPGD models. %Beneficial strain indicated the percentage of nodes on the central pore that experienced the 3 – 10 % tensile strain (first principal strain).

In all pore size groups, the overall tensile strain distribution inside the central pore ranged from 0 to 26%, and the peak of the strain distribution shifted to the right (higher strain) as porosity decreased (**Figure 5b**). The effect of pore parameters on strain distribution in the 3 - 10% strain range was studied, since it was discovered that this range induces anabolism in chondrocytes [31]. Despite the fact that the peaks within 3 – 10% strain range shifted between high and low porosity models, porosity had no effect on the strain distribution pattern in the 250, 500, and 750 groups. Only the 1000 group displayed different strain distribution patterns between high and low porosity models. The 1000-H had the highest peak within the beneficial strain range (3 – 10 %), whereas the 1000-L had the highest peak outside (12%) the beneficial strain range. Additionally, pore size also impacted on strain distribution since increased pore size resulted in increased area with high strain (**Figure 5c**). To better understand the strain distribution inside pores, the percentage of nodes in each model’s central pore that experienced 3 - 10% tensile strain, denoted as %Beneficial strain, was calculated (**Figure 5d**). The findings revealed that, with the exception of the 1000 groups, at least 60% of nodes on the central pore fell within the range of 3 - 10%. As pore size increased and exceeded 500 μm, the level of %Beneficial strain decreased. Porosity also had an impact on the %Beneficial strain because, particularly for the large pore size groups, scaffolds with the same pore size but lower porosity tended to have a higher percentage of beneficial strain. Porous PGD scaffolds with 500 μm pores displayed the highest %Beneficial strain among all the groups.

## 4. Discussion

For cartilage tissue engineering applications, biomaterial scaffolds play a critical role in providing a 3D environment to support cell growth and matrix deposition while protecting the cells from joint loads. Successful scaffold design requires that several essential criteria be met, such as biocompatibility, biodegradability with a favorable resorption rate, suitable pore size and porosity, and mechanical properties that can support tissue growth under native mechanical loads. To develop and model a scaffold that satisfies these requirements, we selected an elastomer, poly (glycerol-dodecanedioate), because it has been shown to be biocompatible, slow-degrading, and nonlinear elastic, and to have shape memory behavior and tailorable mechanical properties, and thus has potential to support cartilage regeneration. To design porous PGD scaffolds to maximize ECM production, the influence of scaffold design parameters, such as pore structure, surface modification, and local strain field, was analyzed. We found that changing the pore size and porosity of the pPGD scaffold altered its surface area, shear modulus, and strain field that developed along the pore surface under load. The type of ligand coating on the PGD surface affected the morphology of hAC in 2-day cultures and influenced both the metabolic activity and ECM production of hAC in 28-day cultures.

In engineered cartilage tissue constructs, the significance of pore diameter and network organization is well-established. Solvent casting, particulate leaching, gas foaming, phase separation, fiber bonding, membrane lamination, melt molding, and freeze drying are examples of common pore-creation techniques [44]. Solvent casting and particulate leaching techniques were developed to more effectively manage the porosity and pore size [45-48]. However, the harsh curing conditions of a thermoset polymer scaffold, e.g., high temperature, limit the possible method for constructing porous structures inside of it. This prevents the use of particulate leaching techniques using organic water-soluble porogens, like sodium alginate, and techniques that only fit thermoplastic polymers. Laser microablation [49], salt leaching [50], and inversely solid freeform fabricated hydroxyapatite mold [13] were successfully conducted on poly (glycerol sebacate) (PGS), a thermoset polymer that has very similar polymerization and crosslinking conditions to PGD, to create uniform internal pores. However, those strategies still have limitations. Laser microablation is constrained by the number of layers and thickness of the porous thermoset scaffold. The limitations of the salt particle leaching method include a narrow achievable range of thickness, uncontrolled interconnectivity, and small pore size of the scaffold. Inversely solid freeform fabrication of a hydroxyapatite mold is a good way to design a customized geometry for thermoset polymer, but it is a complex, multiple-step process. Other methods, such as freezing dry, required the addition of additional materials to achieve pore formation, which altered the native properties of PGS [51]. Our two-step inverse molding method overcomes constraints in sample thickness and complex pore-creation processes.

Our inverse molding technique enables the control of dimension, volume, and organization of 3D porous structures in PGD by arranging wires. However, the achievable pore size with this technique was constrained by the layer-by-layer organization of the wires. The minimum pore size achievable using this method was 250 μm due to the limited resolution of the 3D printed guide and the unstable control of wire arrangement. Previous study has shown that pores < 300 μm in diameter induce chondrocytes to undergo chondrogenic differentiation *in vitro*, while larger pores induce osteogenic differentiation [14]. Other studies show that larger pores (400 – 1000 μm) can improve cell proliferation and cartilage-like matrix deposition [15, 52]. These disparities, which emphasize the significance of pore parameters on chondrocyte function, could be attributed to a variety of material characteristics, manufacturing techniques, and other study-specific scaffold design parameters. Moreover, wire placement allows us to further control the pore size, porosity, and pore arrangement using the inverse molding technique. The wire inverse molding technique was still able to create porous scaffold with a wide range of structural parameters despite having a lower limit on pore size and could therefore be used to optimize chondrocyte behavior during neo-cartilage formation.

Articular cartilage transmits forces across joints, and consequently, the chondrocytes are exposed to a combination of compression, tension, and shear. These mechanical signals are critical regulators of cell behavior and function. It is well known that the magnitude and frequency of the applied local strain cause distinct anabolic or catabolic outcomes in 2D chondrocyte loading experiments *in vitro*. Therefore, the magnitude and distribution of the local strains inside 3D scaffolds under loading are crucial parameters to consider when designing a scaffold. Although the pPGD scaffolds provide a 3D environment for cell proliferation, they also provide 2D surfaces for cell attachment and mechanotransduction due to the large size of the pores created using the inverse molding method. According to the literature, loading chondrocytes with 3 – 10% cyclic tensile strain stimulates their anabolic response in monolayer culture [31]. The positive impact on anabolism was minimal below that range, and higher strains caused catabolic events to predominate [31]. The effects of pore structural parameters on the strain field were investigated using finite element analysis on the pPGD geometry. We found the majority of tensile strains that developed along pore surfaces inside of pPGD scaffolds were at levels shown to be beneficial to chondrocytes, except for the 1000 μm groups (**Figure 5d**). The results in the 1000 μm group could be caused by the large void space in the geometry design step, resulting in a high stress concentration. Studies have shown the large pore of the scaffold not only reduces overall scaffold stiffness, but also has a higher local stress/strain concentration under load than the small pore [53]. Additionally, according to our findings, the proportion of beneficial strain (3 – 10 % for chondrocytes) developed inside pores also depended on porosity (**Figure 5c**). With the same pore size, lower porosity led to a higher percentage of beneficial strain than higher porosity. Therefore, the internal micro-mechanical environment that the cells encounter when loaded in the bioreactor can be impacted by these pore geometries. To optimize the micro-mechanical environment for cells, future research could investigate the combinatorial effects of these parameters on the cell function by coupling the ECM production with the local strain as a first step. Overall, the pPGD scaffold created using our inverse molding process could be used as an *in vitro* model to explore optimal scaffold design parameters for cartilage regeneration.

PGD is a novel biomaterial whose potential as a scaffold for cartilage tissue engineering has never been studied. Our findings demonstrated that PGD, alone or in combination with ECM ligands, promoted chondrocyte survival, attachment, and ECM production (**Figure 4**). It is widely-accepted that the phenotype of chondrocytes may change dramatically when cultured in monolayer for a long period on tissue culture plastic, which is represented by the change of chondrocyte morphology from round to flat and inferior ECM production compared to normal cartilage [54]. Polymeric scaffolds usually provide a stiff 2D environment with a much larger pore sizes than single cell sizes for cell attachment, which may cause a problem of chondrocyte dedifferentiation. Our results revealed that introducing hyaluronic acid ligands on the 2D PGD surface can help to maintain round chondrocyte morphology (**Figure 4**). Previous studies showed the actin cytoskeleton is a major determinant of chondrocyte shape, and the standard actin features of cells such as stress fibrils, filipodia, and lamellipodia were signs for monolayer cultured chondrocytes [42, 43]. Hyaluronic acid-coated PGD retained the rounded morphology of hACs morphology without showing these actin features and maintained high metabolic activity, demonstrating the potential of the microenvironment provided by ligand-coated PGD to preserve chondrocyte phenotype. The cell-ligand binding complex may provide evidence in favor of this. The molecular mechanism underlying the initial adhesion of transplanted chondrocytes to surrounding cartilage in the defect site is the binding of surface receptors to ligands [55]. Integrins on the chondrocytes play a role in the cell proliferation, differentiation, matrix remodeling, and response to mechanical stimuli after initial attachment [19]. Previous study has shown that cell binding to hyaluronic acid, via surface receptors including CD44 and RHAMM, triggers a sophisticated signaling pathway causing chondrocytes to maintain their natural phenotype [56]. Therefore, our results suggested the surface modification involving hyaluronic acid may facilitate the chondrocyte redifferentiation in pPGD scaffolds for cartilage tissue engineering. Furthermore, despite its hydrophobicity, uncoated PGD surface maintained rounder cell shape and induced higher ECM production compared to TCP, indicating its potential for improving cartilage regeneration *in vitro*.

Our results demonstrated the capability of PGD for cartilage regeneration *in vitro*, but more research was necessary to determine whether this material was appropriate to use in a pre-clinical model before using it for specific cartilage applications. The physiochemical or thermomechanical properties of PGD may change during sterilization, medium immersion, and cell culture, due to the nature of hydrolytic degradation in PGD. Studies showed that both *in vitro* and *in vivo* culture environments caused the surface erosion of PGD, which resulted in an increase in transition temperature and stiffness of the polymer [38, 57]. Another study also showed that the sterilization process of PGD by 70% ethanol immersion changed its storage modulus and shape memory properties [58]. Gradually changing the mechanical properties of PGD during culture can result in an alteration in local stress/strain under load, cell phenotype changes, inflammatory responses *in vivo*, and a loss of shape recovery ability at body temperature, limiting the possibility of using this polymer in clinic applications. These results highlight two key directions for engineered PGD scaffolds in soft tissue engineering, 1) understanding the influence of degradation on the mechanical properties of the scaffold and cell behavior during tissue formation, and 2) modifying various scaffold design parameters, such as formulation, surface modification, structural parameters, and so on, to minimize the adverse effect of the material itself and enhance its regenerative function.

## 5. Conclusion

We have developed a scaffold using a novel elastomer that allows for tailoring the 3D pore structure (pore size and porosity), ligand coating, and compressive behavior. This study demonstrates how PGD can be converted into porous scaffolds that facilitate chondrocyte anabolism using an inverse molding technique and surface modification with ECM ligands. The pore size and porosity of porous PGD scaffolds can be tuned by varying the wire diameter or the configuration of wires. HyA coating of the PGD surface was successful in preserving the preferred rounded morphology of hACs during short term culture, and both PGD alone and coated PGD resulted in rising metabolic activity and improved ECM production during long term culture compared to TCP. According to finite element analysis of simulated pPGD structures, the majority of the internal pore surfaces in these scaffolds would experience strain fields that have been shown to be advantageous to chondrocyte function under physiologic loads. Our work demonstrates that this system can be used as a test platform to study how modifications in length scale and organization of the engineered pore network affect cell function. With its aid, we can examine the relationship between pore parameters and other scaffold design factors that also affect cell function, such as ligand presentation and cellular-level strain that develops under loading, to regulate the cellular response to the microenvironment inside engineered scaffolds. Ultimately, the formation of cartilage tissue can be improved by combinatorial tuning of these parameters.

## Acknowledgement

The authors would like to thank Andrea Poli for his support in the mechanical testing, and Xuehui Huang for his help in the confocal imaging.

